# Harnessing photosynthetic bacterium for light-powered biocatalysis

**DOI:** 10.1101/2022.12.20.521182

**Authors:** Yang Zhang, Jifeng Yuan

## Abstract

The traditional whole-cell biocatalysis typically utilizes the heterotrophic microbes as the biocatalyst, which requires carbohydrates to power the cofactor (ATP, NAD(P)H) regeneration. In this study, we sought to harness purple non-sulfur photosynthetic bacterium (PNSB) as the biocatalyst to achieve light-driven cofactor regeneration for cascade biocatalysis. We substantially improved the performance of PNSB-based biocatalysis by using a highly active and conditional expression system, blocking the side-reactions, controlling the feeding strategy, and attenuating the light shading effect. We found that 50 mM ferulic acid could be completely converted to vanillyl alcohol in the recombinant strain, reaching 7.7 g/L vanillyl alcohol. In addition, >99.9% conversion of *p*-coumaric acid to *p*-hydroxybenzoic alcohol (6.21 g/L) was similarly achieved under light-anaerobic conditions. Moreover, we examined the isoprenol utilization pathway (IUP) for pinene synthesis and 13.81 mM pinene (1.88 g/L) with 92.1% conversion rate from isoprenol was obtained. Taken together, these results suggested that PNSB could be a promising host for light-powered biotransformation, which offers an efficient approach for synthesizing value-added chemicals in a green and sustainable manner.

## Main

Biocatalysis plays an essential part of facilitating chemical synthesis in a greener way, as enzymes are usually highly active and operate under mild conditions^1, 2^. The diversity of nature provides a large number of multistep catalytic reactions under the most benign conditions that can be utilized for the production of a variety of chemicals. With the development of synthetic biology, exploiting the enzyme cascades for multistep conversions of renewable feedstocks could afford the relatively toilless admittance to various types of useful value-added compounds^3, 4^. Because of these fascinating features and the dramatic increase in practicable enzymes, cascade enzymatic conversion is turning into a rapid expanding field for chemical synthesis^5^.

Different types of catalysts, such as purified enzymes, immobilized enzymes, cell-free extracts, whole cells, or a mixture of them, can be applied in cascade biocatalysis. A specially-made whole-cell biocatalyst harboring all needed enzymes is regarded as the most economical option among these forms, owing to the advantages as follows^6, 7^: 1) cells are harvested through cultivation at relatively low cost without further downstream purification process; 2) intracellular environment offers cofactor generation to support the reactions; 3) enzymes could be protected with increased life span by cell walls and membranes; 4) co-expression of multiple enzymes within the cells enhances the local concentrations of required enzymes and subsequently decreases the intermediate diffusion in multistep reactions.

In recent years, whole-cell biotransformation has been exploited to convert a variety of substrates such as alkanes, phenols, styrene, fatty acids, ketones, amino acids, and terpene derivatives, into high-value bulk or chiral fine chemicals^7^. The heterotrophic microorganism of *Escherichia coli* represents the most-widely used whole-cell biocatalytic system, and other heterotrophic microorganisms such as *Corynebacterium glutamicum*^8^, *Bacillus subtilis*^9^, and *Pseudomonas putida*^10^, are being established in the recent years. However, all these heterotrophic microorganisms need carbohydrates to support the biomass accumulation and cofactor regeneration, which pose additional costs for their industrial development. For instance, to supply additional ATP and NAD(P)H during the biocatalytic processes^11–14^, additional glucose is added for cofactor regeneration through endogenous cell metabolisms, and in many cases, glucose dehydrogenase (GDH) was supplemented for enhancing NAD(P)H regeneration^13, 15^.

Purple non-sulfur photosynthetic bacterium (PNSB) is widely distributed in nature and it shows extraordinarily versatile metabolism such as photoautotrophic mode using carbon from CO_2_ and energy from light^16^. In particular, the photosynthetic system absorbs light energy to drive electrons transfer along with the generation of a proton gradient across the membrane. Using energy stored in the proton gradient, ATP synthase and NADH-quinone oxidoreductase respectively catalyze the formation of ATP and NADH that are essential for CO_2_ fixation and other metabolic reactions^17^.The ability of utilizing CO_2_ (greenhouse gas) as the carbon source and light energy to support the cell growth and cofactor regeneration makes PNSB a promising and economic whole-cell biocatalyst.

In this study, we attempted to develop a highly-resilient PNSB, namely, *Rhodopseudomonas palustris*, as the platform for light-driven cofactor regeneration to power the biocatalytic processes (Figure 1). Upon systematic optimization of the protein expression system, blocking the side-reactions, controlling the feeding strategy and attenuating the light shading effect, we managed to substantially improve the performance of PNSB-based biocatalysis. We found that vanillyl alcohol (VA), *p*-hydroxybenzyl alcohol (*p*HBA) and pinene were produced near 100% conversion with light-driven cofactor regeneration. Taken together, our work lays a new foundation to harness PNSB as the biocatalyst for chemical productions.

**Figure 1.**
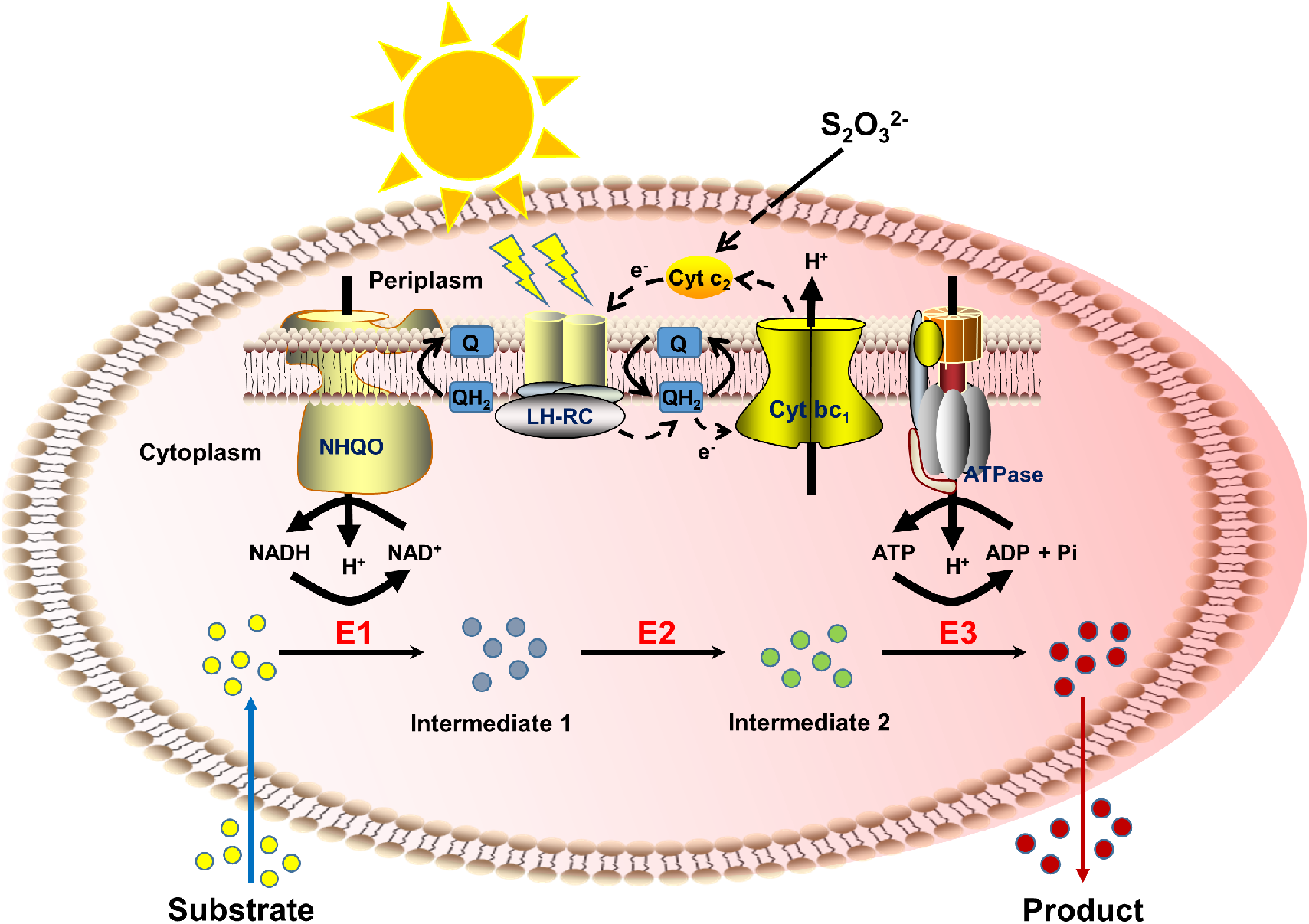
Schematic diagram of light-driven cofactor regeneration for biotransformation using PNSB. Upon excitation by the absorption of light or by energy transfer from light-harvesting complexes, the photosynthetic reaction center reduces ubiquinone (Q) to ubiquinol (QH_2_) using electron (e^-^) from the donor of thiosulfate (S2O_3_^2-^). Cytochrome *bc*_1_ complexes (Cyt *bc*_1_) oxidize QH_2_ and transfer protons (H^+^) across the membrane generating the proton motive force. As a result, ATP and NADH are respectively produced by F_0_F_1_-ATP synthase (ATPase) and NADH: quinone oxidoreductase (NHQO), which can be used in the Calvin cycle to fix CO_2_ into organic molecules. The dotted arrow shows the path of cyclic electron transfer chain from the photosynthetic reaction center to ubiquinone, to Cyt *bc*_1_, to soluble cytochrome *c*_2_ (Cyt *c*_2_). The enzymes (E1^-^E3) employ the cofactors of ATP and NADH generated during anoxygenic photosynthesis to catalyze substrate into the target product. LH^-^RC: light harvesting complexes and photosynthetic reaction center.

## Results

### Development of low-oxygen induced protein expression system under anaerobic conditions

Since the abundance of enzymes is crucial for efficient whole-cell biotransformation, there is a pressing need to develop a highly-active protein expression system in *R. palustris*. In this study, serval promoters to drive the reporter gene of *egfp* were examined in *R. palustris*, including promoters of *puc* operon (P*_puc_*), *puf* operon (P*_puc_*), *bch* operon (P*_bchP_*) and *crt* operon (P*_crtE_*) from *R. palustris*, P_T334-6_ from *Rhodobacter sphaeroides, lac* operon (P*_lac_*) and its variant P*_tac_* from *E. coli*. Specifically, the expression levels of *puc* and *puf* operon encoding the light-harvesting complex II and complex I are relatively high under light-anaerobic conditions; *bch* operon takes charge of the bacteriochlorophyll synthesis and *crt* operon is responsible for the carotenoid synthesis; P_T334-6_, a P*_rsp_7571_*-derived synthetic promoter, showed 32-fold higher activity than that of P*_tac_* under the testing condition in *R. sphaeroides*^18^. As depicted in Figure 2a, five endogenous promoters displayed much higher activity than P*_lac_* but lower than P*_tac_*, and the synthetic promoter of P_T334-6_ exhibited approximately 14-fold higher activity than that of P*_tac_* under light-anaerobic conditions.

**Figure 2.**
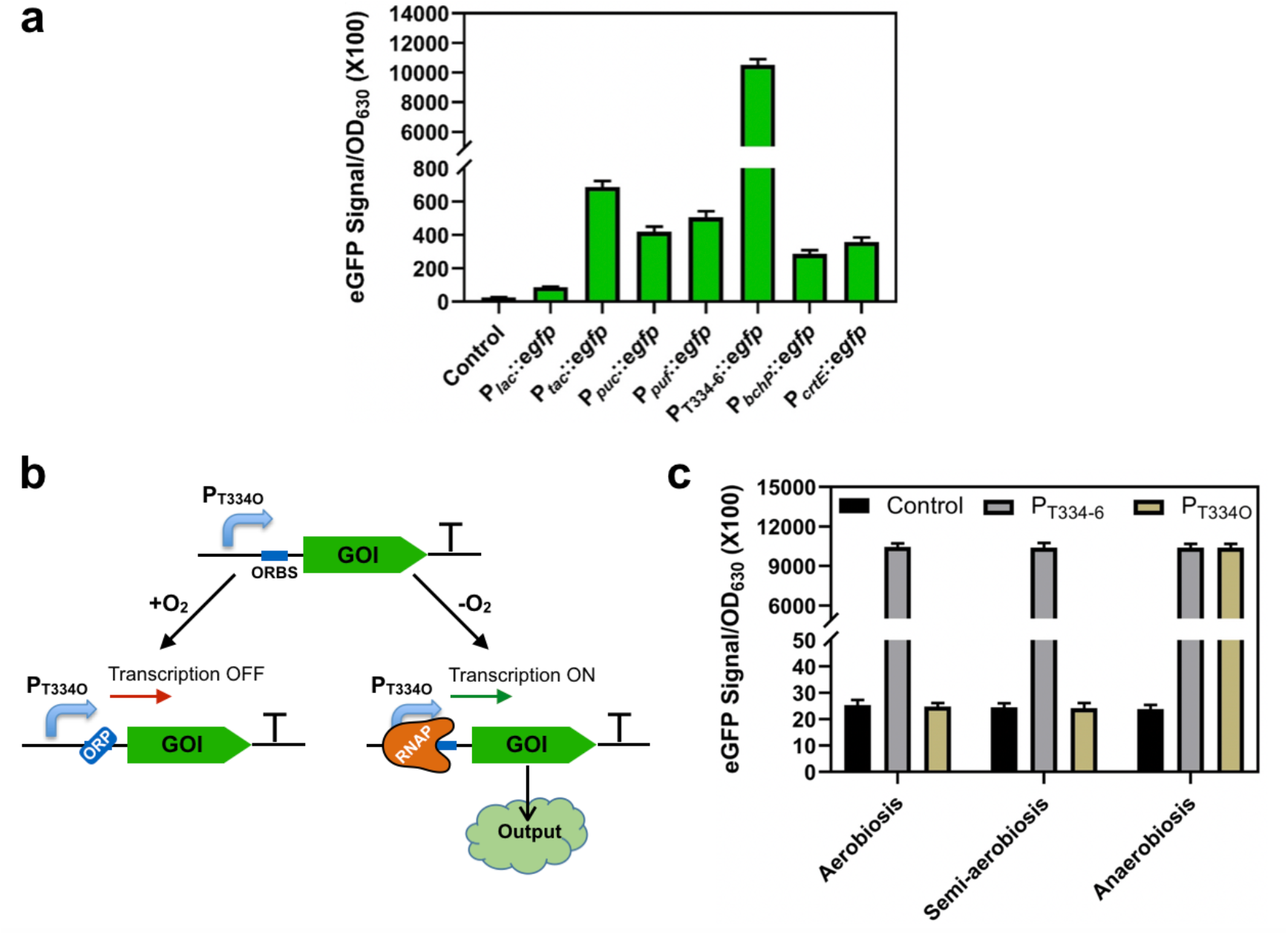
Development of low-oxygen induced protein expression system in *R. palustris*. **a**) The promoter activities based on the reporter gene expression. The promoter activities under light-anaerobic conditions were compared by eGFP fluorescence intensities. The synthetic promoter of P_T334O_, the heterologous promoters of P*_lac_* and P*_tac_* from *E. coli* and the endogenous promoters of P*_puc_*, P*_puf_*, P*_bchP_* and P*_crtE_* were analyzed. **b**) Schematic design of low-oxygen induced protein expression based on the synthetic promoter P_T334O_. ORP: oxygen regular proteins, ORBS: oxygen regular protein binding site, RNAP: RNA polymerase, GOI: genes of interest. **c**) The activities of P_T334-6_ and P_T334O_ were tested at different oxygen levels. The control indicates the strain with the empty plasmid pBRT. All experiments were carried out in triplicate, and the data are presented as mean and standard deviations.

Ideally, the expression of enzymes should be repressed under chemoheterotrophic growth under aerobic conditions, to prevent triggering any protein burden and remain stable maintenance of the genetic construct. According to a previous report, a hybrid promoter of *lacI*^q^-P*_puc_*, was constructed, which is tightly regulated by both isopropyl *β*-D-thiogalactoside (IPTG) and low oxygen level^19^. Due to lack of oxygen-regulatory protein (ORP) binding site information of endogenous P*_puc_* from *R. palustris*, the ORP binding site of P*_puc_* from *R. sphaeroides*^20^ was chosen to insert at the downstream of P_T334-6_ to design P_T334O_ with the aim of using oxygen as the inhibitor (Figure 2b). As shown in Figure 2c, P_T334-6_ activity did not show obvious change at different oxygen levels, indicating that P_T334-6_ itself is insensitive to oxygen tension. Excitingly, we found that ORP of *R. palustris* could interact with ORP binding site from *R. sphaeroides*. As shown in Figure 2c, P_T334O_ exhibited a comparable activity to P_T334-6_ under anaerobic conditions suggesting that the synthetic hybrid promoter of P_T334O_ is tightly activated by low oxygen tension, while P_T334O_ had no obvious activity under aerobic conditions even identical to that under semi-aerobic conditions. Therefore, the synthetic hybrid promoter of P_T334O_ was used for the subsequent cascade biotransformation in *R. palustris*.

### Synthesis of vanillyl alcohol from ferulic acid by cascade biotransformation

Upon the construction of a highly active and controllable protein expression system, we next proceeded to examine PNSB as the biocatalyst for chemical productions. Lignin, the most abundant renewable aromatic biopolymer in nature that makes up 10% to 35% of lignocellulosic biomass, can release diversiform aromatic compounds after depolymerization^21^. Ferulic acid (FA) is one of the common aromatic monomers obtained from lignin depolymerization after the treatment of alkaline oxidation or hydrothermal process^22^. It is well known that *R. palustris* has a robust metabolism to degrade a variety of aromatic compounds, and thus it has a great potential to be a platform for synthesis of high value-added aromatic compounds from renewable lignin-based biomass. To demonstrate the capability of light-driven cofactor regeneration for improved biocatalysis, we proceeded to investigate the biotransformation of FA into vanillyl alcohol (VA), a natural aromatic compound found in several plants such as *Gastrodia elata* Blume and *Vanilla planifolia*, which is known to exhibit a variety of pharmacological activities such as antioxidant, anti-inflammatory, anti-nociceptive, anti-asthmatic, and anti-convulsive activities^23, 24^.

As shown in Figure 3a, the enzymatic cascade contains: FA condenses with coenzyme A (CoA) with expense of ATP by the CoA ligase (encoded by *couB* from *R. palustris*), then FA-CoA undergoes hydration and cleavage to vanillin with the enoyl-CoA hydratase/aldolase (encoded by *couA* from *R. palustris*^25^), and further reduction of vanillin by utilizing the NADH dependent alcohol dehydrogenase encoded by *adh*2 from *Saccharomyces cerevisiae*^26^ to form VA. We therefore constructed two plasmids with modular expression of *couBA* from *R. palustris* and *adh2* from *S. cerevisiae* (Figure S1). Given that CouB and ADH2 require the presence of respective cofactors of ATP and NADH, this biocatalytic route is an excellent choice for investigating the effect of cofactor supply. As depicted in Figure 3b, 2.59 mM VA (51.8%) and 2.41 mM vanillic acid (VAC, 49.2%) were produced from 5 mM FA using strain RVA1 (the wildtype *R. palustris* strain harboring the enzyme cascade of CouBA-ADH2). These findings suggested that PNSB could efficiently intake the substrate and convert FA into vanillin-derived products. However, the accumulation of VAC suggested that the endogenous aldehyde dehydrogenases (ALDHs) could be a limiting factor to achieve efficient VA production in *R. palustris*.

**Figure 3.**
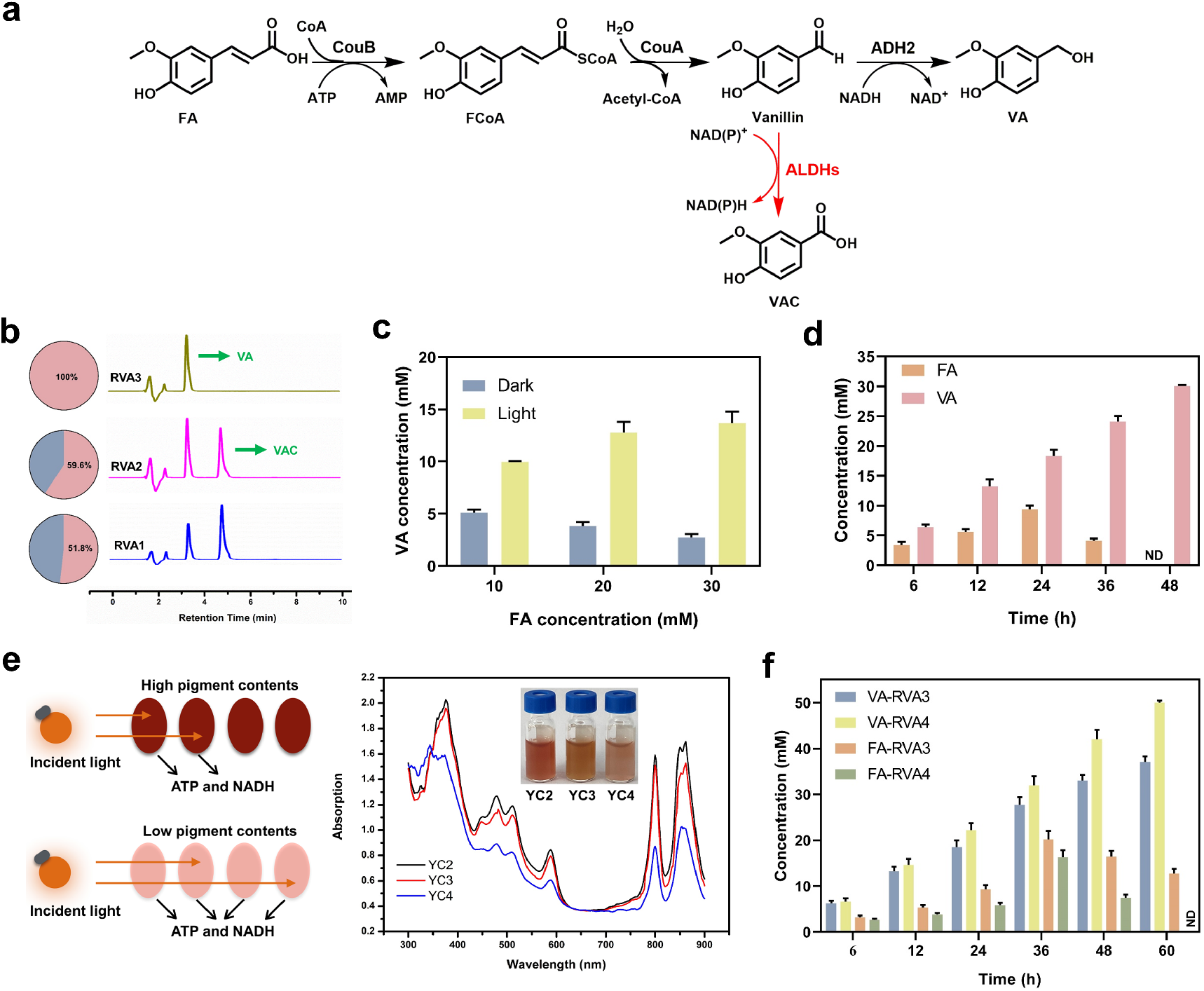
Synthesis of VA from FA by *R. palustris* mediated whole-cell biocatalysis. **a**) Schematic diagram of the biocatalytic route for VA synthesis from FA. FA: ferulic acid, FCoA: ferulyl CoA, VA: vanillyl alcohol, VAC: vanillic acid. **b**) VA productions from 5 mM FA using different recombinant strains. The product of distribution profile and the representative HPLC results are shown. RVA1, RVA2, and RVA3 represent for wild type, the mutant with *RPA1206* deletion, and the mutant with *RPA1206, RPA1687* and *RPA1725* deletions respectively containing CouBA-ADH2. **c**) The effect of light exposure on the VA productions from different concentrations of FA incubating with the resting cells of RVA3 under dark- and light-anaerobic conditions. **d**) Time course of VA production from 30 mM FA by RVA3 using the pulse-feeding approach with supplementing 10 mM FA every 6 h under light-anaerobic conditions. **e**) Manipulation of carotenoid synthesis for reduced light shading. The absorption spectra and colour appearance of *R. palustris* under light-anaerobic conditions are provided. YC3 (*ispA*\*), and YC4 (*ispA*crtE\**) represent the restriction of *ispA* and both *ispA* and *crtE*, respectively. **f**) VA synthesis from 50 mM FA using RVA3 and RVA4 under the pulse-feeding mode. All experiments were conducted in triplicate. Data represent the mean and standard deviations.

### Identification the endogenous factors limiting the vanillyl alcohol synthesis

To avoid the formation of VAC byproduct, we first deleted *RPA1206* (Figure S2a), encoding an ALDH with reported activity on oxidizing *p*-hydroxybenzylaldehyde to *p*-hydroxybenzoic acid under anaerobic conditions^27^. However, the proportion of VA in strain RVA2 (strain YC1 harboring CouBA-ADH2) was slightly improved from 51.8% to 59.6%, suggesting that there are other ALDHs involved in VAC formation. Next, we proceeded to delete *RPA1687* (coniferyl aldehyde reductase) and *RPA1725* (aldehyde reductase to phenolic acid) to further disrupt VAC formation from vanillin (Figure S2a). Encouragingly, 5 mM VA with ~100% conversion was obtained in *R. palustris* RVA3 (strain YC2 harboring CouBA-ADH2) and no VAC was detected (Figure 3b). In addition, there was no obvious differences of cell growth and light absorption between the wild type and the mutant strain with triple deletion of *RPA1206*, *RPA1687* and *RPA1725* (Figure S2b and S2c).

To confirm the essentiality of light to power the biocatalysis, we next compared the VA production between dark- and light-anaerobic conditions. As depicted in Figure 3c, we found that the conversions of FA to VA were much higher under light-anaerobic conditions when compared to those under dark conditions. For instance, 13.68 mM VA were obtained with 45.6% conversion under light-anaerobic conditions, whereas only 9.1% conversion were reached under dark-anaerobic conditions. Therefore, we concluded that light plays an indispensable role in regenerating the cofactors to accelerate the biotransformation. Since the biocatalytic system experienced a severe substrate inhibition, we next proceeded the pulse-feeding strategy by periodically adding 10 mM FA every 6 h and 30 mM FA was fully converted into VA within 48 h (Figure 3d).

### Attenuating the light shading effect for improved biocatalytic efficiency

Since whole-cell biotransformation typically uses high-cell density for the catalysis, the pigment content in *R. palustris* would cause the shading effect that prevents the transmission ability of light, thereby diminishes the cofactor regeneration (Figure 3e). In *R. palustris*, the main pigment is attributed to the carotenoid content, which is synthesized from the methylerythritol phosphate (MEP) pathway. Given farnesyl pyrophosphate synthase (FPPS encoded by *ispA*) and geranylgeranyl diphosphate synthase (GGPPS encoded by *crtE*) are involved in carotenoids synthesis, the promoter P*_lac_* with a relatively low activity (Figure 2b) was used to replace the promoters of *ispA* and *crtE* to obtain *R. palustris* YC3 (*ispA*\*) and YC4 (*ispA* crtE\**) based on the control stain YC2 as shown in Figure S3a. The decrease of *ispA* expression resulted in a slight lower light absorption compared to that of the control due to the presence of *crtE* while the decreases of *ispA-crtE* expressions gave rise to the remarkable lower light absorption (Figure 3e), implying that the pigment biosynthesis were significantly weakened. Besides, the colour appearance of strains correlated well with the light absorption profile (Figure 3e). Notably, we did not observe noticeable change of the growth of all the recombinant strains (Figure S3b), suggesting that the above-mentioned modifications did not obviously alter the photosynthetic ability.

When the engineered strains with less light shading were used for the biocatalytic formation of VA from FA, the overall catalytic efficiencies were substantially improved in these strains. As shown in Figure 3f, strain RVA4 (YC4 harboring CouBA-ADH2) displayed a better performance at high FA concentration, and 50 mM FA was converted to 7.7 g/L VA with > 99.9% conversion within 60 h, whereas only 74.1% conversion was obtained by the control of RVA3. Therefore, we have further improved the performance of PNSB-mediated biocatalyst by attenuating the light shading, which would be of significant interest for the future industrial applications when the light transmission becomes a bottleneck.

### Synthesis of *p*-hydroxybenzyl alcohol from *p*-coumaric acid by cascade biotransformation

To demonstrate the broad utility of CouBA-ADH2, we also used the same enzyme cascade to produce *p*-hydroxybenzyl alcohol (*p*HBA) from *p*-coumaric acid (*p*CA), another abundant lignin-derived compound (Figure S4a). When *p*CA concentration increased from 5 to 30 mM, 5.0 to 15.11 mM *p*HBA were gained by RHBA1 (as same as the strain RVA3) along with substrate conversion descending from 100% to 50.4% under light-anaerobic conditions, whereas low titers of 3.52 to 3.17 mM *p*HBA (70.4% to 10.6% conversion) were synthesized under dark conditions as described in Figure S4b. Moreover, no *p*-hydroxybenzoic acid (*p*HBAC) was observed during the biocatalysis based on the HPLC result (Figure S5). We further proceeded the pulse-feeding strategy to reduce substrate inhibition and 3.72 g/L *p*HBA with about 100% conversion from 30 mM *p*CA was achieved in 48 h (Figure S4c). In addition, we also examined the engineered strain with attenuated photosynthetic pigments and more efficient *p*HBA synthesis was obtained in strain RHBA2 (as same as the strain RVA4) at high *p*CA concentration of 50 mM. Specially, the substrate of *p*CA was fully transformed into 6.21 g/L *p*HBA within 60 h, while only 4.73 g/L *p*HBA with 76.2% conversion was reached by the control and 10.85 mM substrate remained (Figure S4d). Therefore, we concluded that the enzyme cascade of CouBA-ADH2 could be applied for synthesizing other aromatic alcohols by simply varying lignin-derived monomers.

### Synthesis of pinene from isoprenol by cascade biotransformation

To further expand the applicability of light-powered biocatalysis, we also examined isoprenol utilization pathway (IUP) based terpene synthesis. In this study, we used pinene synthesis from isoprenol as an example, which requires large amounts of ATP for substrate phosphorylation (Figure 4a). Cascade 1: isoprenol undergoes two-step phosphorylation to isopentenyl diphosphate (IPP) by using a *E. coli* kinase (ThiM)^28^ and isopentenyl phosphate kinase (IPK) from *Arabidopsis thaliana*^29^, followed by interconversion of IPP and dimethylallyl diphosphate (DMAPP) using isopentenyl pyrophosphate isomerase (Idi) from *E. coli*. Cascade 2: condensation of IPP and DMAPP by geranyl diphosphate synthase (GPPS) from *Abies grandis* gives GPP which serves as the substrate for pinene synthesis via the further action by pinene synthase (PS) from *A. grandis*^30^. To mitigate the competing pathways of host metabolism, we used a direct fusion of GGPS with PS as previously reported^30^.

**Figure 4.**
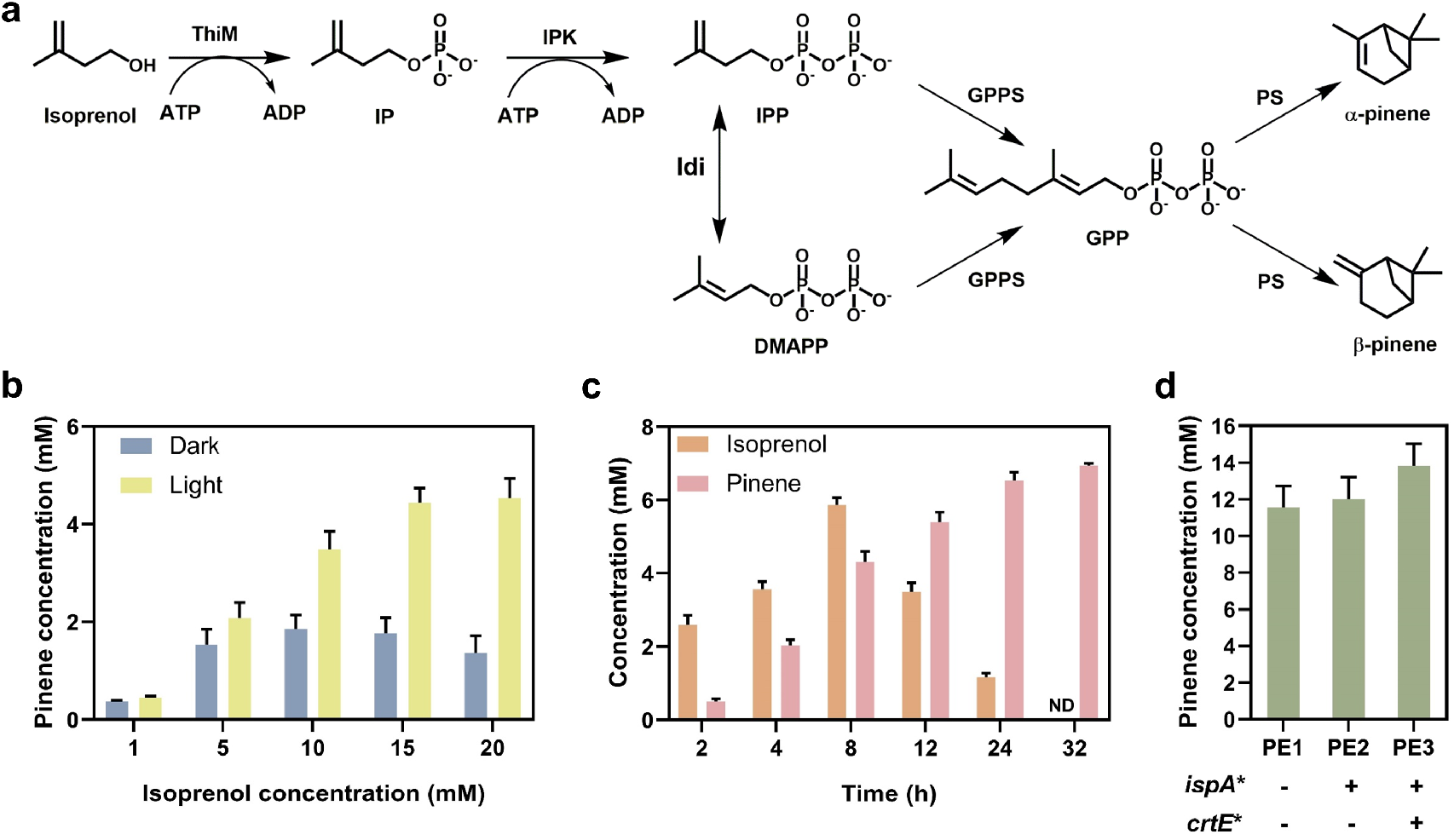
Synthesis of pinene from isoprenol by *R. palustris* mediated whole-cell biocatalysis. **a**) Schematic diagram of IUP-coupled pinene synthesis from isoprenol. **b**) Biotransformation of isoprenol into pinene under dark- and light-anaerobic conditions by PE1. **c**) Time course of the pinene production from 20 mM isoprenol by PE1 using the pulse-feeding approach under light-anaerobic conditions. **d**) Biotransformation of 30 mM isoprenol into pinene by the engineered strains with reduced light shading using the pulse-feeding approach under light-anaerobic conditions. PE1, PE2, and PE3 represents for wild type, YC5, and YC6 harboring the cascades ThiM-IPK-Idi-GPPSps, respectively. All experiments were conducted in triplicate. Data represent the mean and standard deviations.

The *R. palustris* containing the desired enzyme cascades (Figure S6) was then tested for pinene synthesis from isoprenol in a two-phase system containing 20% *n*-dodecane. As depicted in Figure 4b, the yields of pinene under light-anaerobic conditions reached 90%-45.4%, which were significantly higher than those under dark conditions (74%-13.6%). These results suggested that the light-driven ATP regeneration in *R. palustris* is still effective in improving the overall conversion of isoprenol to pinene. Since the fusion proteins of GPPS-PS could create a direct channel between GPP and pinene that facilitates the flux to product (pinene) and alleviates the GPP inhibition to PS^31^, and two-phase system could alleviate the pinene toxicity and its inhibition to PS, we reasoned that the low titers of pinene biosynthesis might be caused by the toxicity of isoprenol to the host or substrate inhibition of isoprenol. We next proceeded with periodical addition of 4 mM isoprenol every 2 h until the final content reached 20 mM by utilizing pulse-feeding strategy. We were excited to find that 6.94 mM pinene was achieved and the conversion was substantially enhanced from 45.4% of direct feeding to 69.4% (Figure 4c), and no isoprenol was detected after 32 h (Figure S7). Furthermore, the engineered strains with less light absorption were also examined for pinene synthesis from 30 mM isoprenol. As shown in Figure 4d, we found the engineered strain with reduced light shading could substantially improve the overall conversion of isoprenol to pinene, reaching 13.81 mM pinene (1.88 g/L) in YC6 derivative of PE3 with 92.1% conversion.

## Discussion

Biocatalysis is considered to be a sustainable and environment-friendly alternative to chemical synthesis. In this study, we have developed *R. palustris* as the whole-cell biocatalyst, and harnessed the light to power cofactor regeneration for biotransformation applications. The performance of PNSB-based biocatalysis was substantially improved by using a highly active and conditional expression system, blocking the side-reactions, controlling the feeding strategy and attenuating the light shading effect. When compared to the traditional heterotrophic microorganisms that require additional carbohydrates for cofactor regeneration, photoautotrophic PNSB-based biocatalyst might avoid competition with food industry and significantly reduce the operating cost.

In this study, we have demonstrated light-powered biotransformation for VA and *p*HBA productions using the endogenous CoA-dependent non-β-oxidation route of *R. palustris* together with NADH-dependent ADH2 from *S. cerevisiae*. Both FA and *p*CA were efficiently converted to the corresponding “Cn-2” alcohols, with ~100% conversion under light-anaerobic conditions. When compared to previous studies^32, 33^, we have reached better titers of VA (7.7 g/L) and *p*HBA (6.21 g/L), suggesting that PNSB is an appealing and feasible host for light-powered lignin valorization. In the future, it will be possible to engineer *R. palustris* to synthesize diverse aromatic compounds such as vanillylamine^34^, protocatechuic acid and gallic acid^12^ in a similar manner. In addition, we also developed a biocatalytic system for pinene synthesis from isoprenol and 9.21 mM pinene (1.25 g/L) with 92.1% conversion was achieved, which was much higher than the titers obtained by traditional metabolic engineering efforts^35,36^. For the future production of other terpenes such as sesquiterpene, it will require a proper balance of DMAPP and IPP levels by adjusting Idi expression or simply use a fixed ratio of prenol and isoprenol without introducing Idi-mediated isomerization to achieve the theoretical maximum.

To further improve light-powered biotransformation, we also engineered the carotenoid biosynthetic pathway of *R. palustris* to weaken the light shading effect and improve the transmission of light through the cell culture. Our results confirmed that the engineered strains with lower light absorption could accelerate the conversion process. It was reported that disruption of *puc* operon^37^ or overexpression of *pufQ*^38^ (a regulatory gene to *puf* operon), could decrease the pigment contents with low light absorption, which might be conducive to ATP and NADH synthesis. Due to photophosphorylation coupling with electron transport during photosynthesis, slight overexpression of *cycA* encoding for cytochrome *c*2, an important electron carrier in electron transport chain, was found to improve ATP and NADH synthesis^39^. In addition, transposon mutagenesis screening has identified several mutants with reduced pigments^40, 41^. These engineering strategies might be similarly implemented in our light-powered biotransformation to further improve the strain performance in the future work.

Although we have achieved impressive results based on PNSB as the biocatalyst, there are several limitations to be considered. Firstly, the product or substrate should not interfere the performance of photosynthetic apparatus, otherwise it will decrease the cofactor regeneration. Secondly, anoxygenic photosynthesis of PNSB could restrict its applications in some redox reactions that require O_2_. Autotrophic microorganisms such as cyanobacteria have been applied for direct photosynthetic recycling of CO_2_ into value-added products^42, 43^, but there are limited reports on utilizing cyanobacteria for whole-cell biocatalysis. Considering cyanobacteria afford encouraging source of reducing power of NADPH, it will be promising to develop such platform for oxygenic biocatalysis applications. Therefore, more efforts are needed to expand this research area and develop the applications of autotrophic microbes to synthesize chemicals by whole-cell biocatalysis.

## Methods

### Strains and cultivation

*R. palustris* CGA009 was employed as the base strain to engineer the chassis cell for biotransformation. For chemoheterotrophic growth, *R. palustris* was cultivated in Sistrom’s mineral medium (MedS)^44^ at 35°C under the aerobic condition supplemented with antibiotics (10 μg/mL kanamycin, and 12 μg/mL gentamycin) when needed. For photoautotrophic growth, *R. palustris* was cultivated in modified Sistrom’s mineral medium (MedC) in which succinate was replaced by 10 mM NaHCO_3_ and 1 mM Na_2_S_2_O_3_ at 35°C under the light-anaerobic conditions (4000 lux) with 50 kPa of 80% H_2_/20% CO_2_^45^ supplemented with antibiotics if necessary. *E. coli* S17-1 was applied as the competent cell for plasmid construction and cultivated in Luria-Bertani (LB) medium at 37°C supplemented with antibiotics (50 μg/mL kanamycin, and 12 μg/mL gentamycin) when needed. All strains used in this study are listed in Table S1.

### Plasmids construction

PCR amplification of all genes was performed with High Fidelity Phusion DNA polymerase or Taq polymerase from New England Biolab (Ipswich, MA, USA). All restriction enzymes and T4 DNA ligase were obtained from New England Biolab (Ipswich, MA, USA). The rrnB T1 terminator was synthesized by GenScript (Nanjing, China) cut with *Xho*I and *Sal*I, then cloned into pBdRSf^46^ via *Xho*I to give pBRT confirmed by sequencing. The untranslated regions of approximately 500 bp upstream from *puc*, *puf, bch* and *crt* operons were amplified from the genomic DNA of *R. palustris* CGA009 with primers Ppuc-F/Ppuc-R, Ppuf-F/Ppuf-R, PbchP-F/PbchP-R, and PcrtE-F/PcrtE-R, and then cloned into pBRT between *Eco*RI and *Kpn*I sites to obtain pBRPpuc, pBRPpuf, pBRPbchP, and pBRPcrtE, respectively. The P_T334-6_ promoter^18^ was synthesized by GenScript (Nanjing, China) then cloned into pBRT between *Eco*RI and *Kpn*I sites to yield pBRPt334-6, and the DNA fragments containing the P_T334-6_ promoter and the oxygen-regulatory protein binding site of P*_puc_* from *R. sphaeroides* attained by PCR with primers Pt334O-F/Pt334O-R was digested with *Eco*RI and *Kpn*I and then cloned into pBRT to produce pBRPt334O. Besides, P*_lac_* and P*_tac_* promoters amplified from the vector pBBR1MCS-2 and pBBR-tacGFP^30^ with primers Plac-F/Plac-R and Ptac-F/Ptac-R were respectively cloned into pBRT digested by *Eco*RI and *Kpn*I to produce pBRPlac and pBRPtac. Finally, the *egfp* fragment was cloned into these plasmids via *Bam*HI and *Xho*I to give pBRpuc-eGFP, pBRpuf-eGFP, pBRbchP-eGFP, pBRcrtE-eGFP, pBRt334-6-eGFP, pBRt334O-eGFP, pBRlac-eGFP, and pBRtac-eGFP, respectively.

To replace the kanamycin resistance marker of pBRPt334O to gentamycin resistance, the DNA fragment of gentamycin resistance was amplified from plasmid pZJD29c with primers Gen-F/Gen-R, and then circular PCR was used to create pGenPt334O with gentamycin resistance. To obtain pBRT334O-CouBA, CoA ligase gene (*couB*)^27^ and enoyl-CoA hydratase/aldolase gene (*couA*)^27^ were respectively amplified from *R. palustris* genome using primers couB-F/couB-R and couA-F/couA-R, and then cloned together into pBRPt334O between *Bam*HI and *Xho*I sites. To acquire pGenT334O-ADH2, alcohol dehydrogenase 2 (ADH2 encoded by *adh2*) gene from *S. cerevisiae*^47^ amplified with primers ADH2-F/ADH2-R was cloned into pGenPt334O between *Bam*HI and *Xho*I sites. The fragments of hydroxyethylthiazole kinase (ThiM) gene from *E. coli* amplified with primers ThiM-F/ThiM-R, isopentenyl phosphate kinase (IPK) gene from *A. thaliana* amplified with primers IPK-F/IPK-R, and isopentenyl pyrophosphate isomerase (Idi) gene from *E. coli* amplified with primers Idi-F/Idi-R were cloned together into pBRPt334O digested with *Bam*HI and *Xho*I to give pBRT334O-ThiM-IPK-Idi. The fragments of GPPS-PS fusion amplified from pBBR-αGppsPs^30^ with primers GPPSps-F1/GPPSps-R1 and GPPSps-F2/GPPSps-R2 were cloned into pGenPt334O digested with *Bam*HI and *Xho*I to give pGenT334O-GPPSps. All oligonucleotides synthesized from GenScript (Nanjing, China) and plasmids used in this study are listed in Table S2 and S3, respectively.

### Di-parental conjugation

Conjugation mating was implemented for transforming plasmids into *R. palustris*. *E. coli* S17-1 bearing the desired plasmids and *R. palustris* were employed as donor strain and receptor strain, respectively. After *R. palustris* was aerobically cultivated in 10 mL MedS at 35°C for 24 h and *E. coli* S17-1 was cultivated in 5 mL LB at 37°C for overnight, the cells were harvested and washed once with fresh MedS, and resuspended with 1 mL MedS. Then *E. coli* and *R. palustris* were mixed as a ratio of 3:10 (V/V), and the mixture was transferred on a MedS agar plate incubating at 35°C for 24 h. The cultures were harvested, washed once with fresh MedS, resuspended with 800 μL MedS, and spread on a MedS agar plate with appropriate antibiotics. Colonies typically appeared after 2-3 days at 35°C.

### Genome editing in *R. palustris*

The suicide plasmid pZJD29c^48^ mediated homologous recombination was used for genome editing in *R. palustris*. To attain pZJ-ΔALDH1 for *RPA1206* deletion, the upstream and downstream fragments of the *aldh1* gene were amplified from CGA009 genome with respective primers ALDH1-F1/ALDH1-OE-R1 and ALDH1-OE-F2/ALDH1-R2 assembled by overlapping PCR, then cloned into the *Sac*I and *Kpn*I sites on pZJD29c. The similar way was used to generate pZJ-ΔALDH_2_ for *RPA1687* deletion and pZJ-ΔALDH3 for *RPA1725* deletion. In order to implement the promoter replacement, the upstream and downstream fragments of the *ispA* promoter amplified from *R. palustris* CGA009 genome with respective primers ispA-uF/ispA-OE-uR and ispA-OE-dF/ispA-dR and the fragment of P*_lac_* promoter amplified from pBBR1MCS-2 with primers Plac-OE-iF/Plac-OE-iR were assembled by overlapping PCR and cloned into pZJD29c cut with *Xba*I and *Kpn*I to generate pZJ-Plac-ispA. Based on the similar method, pZJ-Plac-crtE was attained using primers crtE-uF/crtE-OE-uR, Plac-OE-cF/Plac-OE-cR, and crtE-OE-dF/crtE-dR. Single colonies harboring pZJD29c-derived plasmids were picked and cultivated in 1 mL MedS for overnight, and then screened on MedS plate containing 10% sucrose until colonies appeared. Positive mutants were confirmed by diagnostic PCR and DNA sequencing.

### Fluorescence intensity assays

To measure the eGFP fluorescence, *R. palustris* recombinants were aerobically cultivated in 5 mL MedS at 35°C until OD_630_ (optical density at 630 nm) reached approximately 0.6, then the culture was cultivated at 30°C under light-anaerobic conditions for 2 days. OD_630_ values and fluorescence intensities were analyzed with a microplate reader, Synergy H1 (BioTek, USA). The excitation and emission of eGFP were set at 485 nm and 510 nm respectively, and the fluorescence intensities were normalized to OD_630_.

### RNA preparation and real-time qPCR

Total RNA of *R. palustris* was extracted with the Bacterial RNA Kit (Omega Bio-Tek, GA, USA) based on the protocol and treated with DNase I to remove the genomic DNA. Then, cDNA synthesis was conducted with M-MuLV First Strand cDNA Synthesis Kit (Sangon Biotech, China) with Oligo(dT) as primers according to the instruction. Realtime qPCR was carried out using Perfectstart SYBR Green qPCR Master Mix kit (Omega Bio-Tek, GA, USA) with a thermal cycler (Agilent AriaMx, Agilent, USA) and the corresponding primers are presented in Table S2. The expression levels of genes were analyzed by relative quantification-ΔΔ*Ct* method in which *Ct* values were determined from house-keeping gene (16S rRNA) and target genes in test strains.

### Biocatalysis procedures

The recombinant strains of *R. palustris* were aerobically incubated in 5 mL MedS at 35°C and 250 rpm for overnight. Then the fresh culture was transferred into a 250 mL shake flask containing 100 mL MedC and aerobically grown at 35°C until OD_630_ attained approximately 0.6. Afterwards, the culture was incubated at 30°C under photoautotrophic conditions facilitating for protein expression.

After reaching an OD_630_ of 1.8-2.0, the cells were harvested by centrifugation at 7000 rpm, 4°C for 10 min and washed once with ice-cold water and once with KP buffer (200 mM, K_2_HPO_4_·3H_2_O 42.90 g/L, KH_2_PO_4_ 1.63 g/L, pH 7.0), then resuspended in KP buffer to a cell density of 50 g-cdw/L. For biotransformation, 2 mL reaction mixture generally comprised of 15 g-cdw/L cell suspension, a certain amount of substrate, 10 mM NaHCO_3_, 1 mM Na_2_S_2_O_3_ and KP buffer (200 mM, pH 7.0). Na_2_S_2_O_3_ and NaHCO_3_ were added to the reaction mixture for the purpose of cofactor regeneration *via* cellular metabolism. Besides, 5 mM MgCl_2_ and 1 mM MnCl_2_ were supplemented for pinene synthesis. All biotransformations were carried out at 30°C and 250 rpm under dark- and light-anaerobic conditions.

### Analytical methods

FA, *p*CA, *p*HBA, *p*HBAC, VA, and VAC in the aqueous phase were analyzed by a high-performance liquid chromatography (HPLC, Shimadzu LC-20A, Japan) equipped with a photodiode array detector and a reversed-phase column (C18, 150 mm × 4.6 mm × 5 μm) under 40°C. Mobile phase: 70% ultrapure H_2_O with 0.1% trifluoroacetic acid and 30% acetonitrile. Flow rate: 1 mL/min. Detection wavelength: 210 nm. Isoprenol and pinene in the organic phase were measured by a gas chromatography (GC, Shimadzu GC-2030, Japan) equipped with a flame ionization detector and a Rtx-5 column (30 m × 0.25 mm × 0.25 μm). Nitrogen was used as a carrier with a flow rate of 1.0 mL/min. The column temperature was first kept at 80°C for 2 min, then increased to 190°C at a rate of 5°C/min, and finally increased to 300°C by 20°C/min. External authentic standards were used for plotting the standard curve. The product and substrate levels were calculated accordingly.

## Acknowledgments

We acknowledge financial support from Xiamen University under grant no. 0660X2123310, the Natural Science Foundation of Fujian Province of China no. 2020J05011, Guangdong Basic and Applied Basic Research Foundation no. 2021A1515110340, Daan Gene (20223160A0063) and ZhenSheng Biotech. The authors would also like to thank Lu Zhang, Junyi Wang and Prof. Mingfeng Cao for their assistance during the preparation of manuscript.

## Author Contributions

J.Y. conceived the project. Y.Z. and J.Y. designed the experiments. Y.Z. carried out the experiments and collected the data. Y.Z. and J.Y. wrote the manuscript.

## TOC

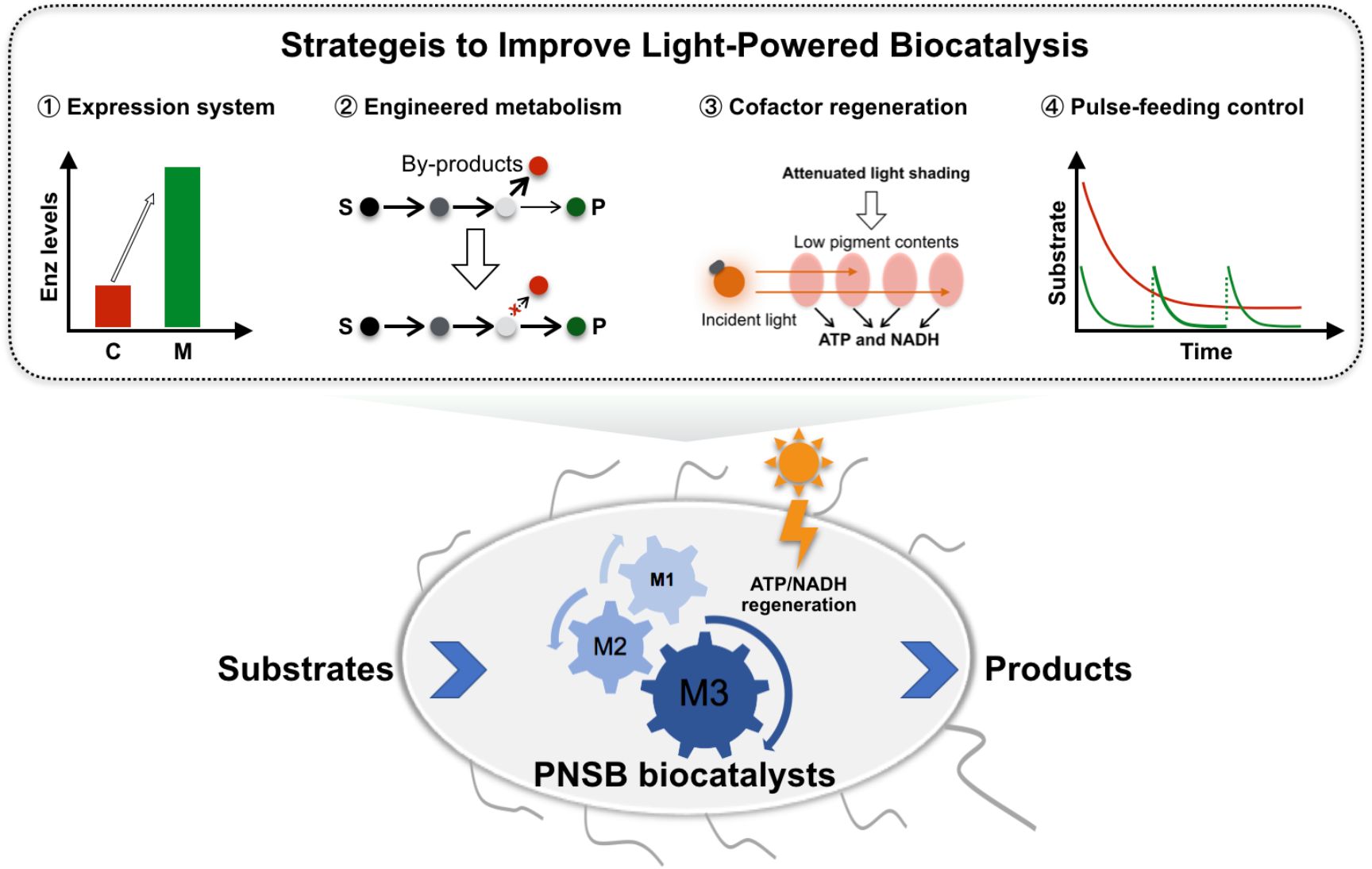

## References

1. Bornscheuer, U.T. et al. Engineering the third wave of biocatalysis. Nature 485, 185–194 (2012).

2. Reetz, M.T. Biocatalysis in organic chemistry and biotechnology: Past, present, and future. J. Am. Chem. Soc. 135, 12480–12496 (2013).

3. Kohler, V. & Turner, N.J. Artificial concurrent catalytic processes involving enzymes. Chem. Commun. 51, 450–464 (2015).

4. Muschiol, J. et al. Cascade catalysis-strategies and challenges *en route* to preparative synthetic biology. Chem. Commun. 51, 5798–5811 (2015).

5. Schrittwieser, J.H., Velikogne, S., Hall, M. & Kroutil, W. Artificial biocatalytic linear cascades for preparation of organic molecules. Chem. Rev. 118, 270–348 (2018).

6. Wachtmeister, J. & Rother, D. Recent advances in whole cell biocatalysis techniques bridging from investigative to industrial scale. Curr. Opin. Biotechnol. 42, 169–177 (2016).

7. Wu, S. & Li, Z. Whole-cell cascade biotransformations for one-pot multistep organic synthesis. ChemCatChem 10, 2164–2178 (2018).

8. Son, J. et al. Production of *trans*-cinnamic acid by whole-cell bioconversion from L-phenylalanine in engineered *Corynebacterium glutamicum*. Microb. Cell Fact. 20, 145 (2021).

9. Hossain, G.S. et al. Improved production of α-ketoglutaric acid (α-KG) by a Bacillus subtilis whole-cell biocatalyst via engineering of L-amino acid deaminase and deletion of the α-KG utilization pathway. J. Biotechnol. 187, 71–77 (2014).

10. Upadhyay, P., Singh, N.K., Tupe, R., Odenath, A. & Lali, A. Biotransformation of corn bran derived ferulic acid to vanillic acid using engineered *Pseudomonas putida* KT2440. Prep. Biochem. Biotechno. 50, 341–348 (2020).

11. France, S.P. et al. One-pot cascade synthesis of mono- and disubstituted piperidines and pyrrolidines using carboxylic acid reductase (CAR), ω-transaminase (ω-TA), and imine reductase (IRED) biocatalysts. ACS Catal. 6, 3753–3759 (2016).

12. Fu, B., Xiao, G., Zhang, Y. & Yuan, J. One-pot bioconversion of lignin-derived substrates into gallic acid. J. Agric. Food Chem. 69, 11336–11341 (2021).

13. Ge, J., Yang, X., Yu, H. & Ye, L. High--yield whole cell biosynthesis of Nylon 12 monomer with self-sufficient supply of multiple cofactors. Metab. Eng. 62, 172–185 (2020).

14. Wu, S., Liu, J. & Li, Z. Biocatalytic formal anti-Markovnikov hydroamination and hydration of aryl alkenes. ACS Catal. 7, 5225–5233 (2017).

15. Wang, P., Yang, X., Lin, B., Huang, J. & Tao, Y. Cofactor self-sufficient wholecell biocatalysts for the production of 2-phenylethanol. Metab. Eng. 44, 143–149 (2017).

16. Larimer, F.W. et al. Complete genome sequence of the metabolically versatile photosynthetic bacterium *Rhodopseudomonas palustris*. Nat. Biotechnol. 22, 55–61 (2004).

17. Johnson, E.T. & Schmidt-Dannert, C. Light-energy conversion in engineered microorganisms. Trends Biotechnol. 26, 682–689 (2008).

18. Shi, T. et al. Screening and engineering of high-activity promoter elements through transcriptomics and red fluorescent protein visualization in *Rhodobacter sphaeroides*. Synth. Syst. Biotechnol. 6, 335–342 (2021).

19. Hu, Z., Zhao, Z., Pan, Y., Tu, Y. & Chen, G. A powerful hybrid *puc* operon promoter tightly regulated by both IPTG and low oxygen level. Biochem-Moscow 75, 519–525 (2010).

20. McGlynn, P. & Hunter, C.N. Isolation and characterization of a putative transcription factor involved in the regulation of the *Rhodobacter sphaeroides pucBA* operon. J. Biol. Chem. 267, 11098–11103 (1992).

21. Cai, C., Xu, Z., Zhou, H., Chen, S. & Jin, M. Valorization of lignin components into gallate by integrated biological hydroxylation, O-demethylation, and aryl side-chain oxidation. Sci. Adv. 7, eabg4585 (2021).

22. Santos, J.H., Martins, M., Silvestre, A.J., Coutinho, J.A. & Ventura, S.P.J.G.C. Fractionation of phenolic compounds from lignin depolymerisation using polymeric aqueous biphasic systems with ionic surfactants as electrolytes. Green Chem. 18, 5569–5579 (2016).

23. Jung, H.J., Song, Y.S., Lim, C.J. & Park, E.H. Anti-angiogenic, antiinflammatory and anti-nociceptive activities of vanillyl alcohol. Arch. Pharm. Res. 31, 1275–1279 (2008).

24. Hsieh, C.L. et al. Anticonvulsive and free radical scavenging activities of vanillyl alcohol in ferric chloride-induced epileptic seizures in Sprague-Dawley rats. Life Sci. 67, 1185–1195 (2000).

25. Hirakawa, H., Schaefer, A.L., Greenberg, E.P. & Harwood, C.S. Anaerobic *p*-coumarate degradation by *Rhodopseudomonas palustris* and identification of CouR, a MarR repressor protein that binds *p*-coumaroyl coenzyme A. J. Bacteriol. 194, 1960–1967 (2012).

26. Chen, Y. et al. High-yielding protocatechuic acid synthesis from L-tyrosine in *Escherichia coli*. ACS Sustain. Chem. Eng. 8, 14949–14954 (2020).

27. Pan, C. et al. Characterization of anaerobic catabolism of *p*-coumarate in *Rhodopseudomonas palustris* by integrating transcriptomics and quantitative proteomics. Mol. Cell. Proteomics 7, 938–948 (2008).

28. Clomburg, J.M., Qian, S., Tan, Z., Cheong, S. & Gonzalez, R. The isoprenoid alcohol pathway, a synthetic route for isoprenoid biosynthesis. Proc. Natl. Acad. Sci. U. S. A. 116, 12810–12815 (2019).

29. Chatzivasileiou, A.O., Ward, V., Edgar, S.M. & Stephanopoulos, G. Two-step pathway for isoprenoid synthesis. Proc. Natl. Acad. Sci. U. S. A. 116, 506–511 (2019).

30. Wu, X. et al. Biosynthesis of pinene in purple non-sulfur photosynthetic bacteria. Microb. Cell Fact. 20, 101 (2021).

31. Sarria, S., Wong, B., Martín, H.G., Keasling, J.D. & Peralta-Yahya, P. Microbial synthesis of pinene. ACS Synth. Biol. 3, 466–475 (2014).

32. Chen, Z. et al. Establishing an artificial pathway for *de novo* biosynthesis of vanillyl alcohol in *Escherichia coli*. ACS Synth. Biol. 6, 1784–1792 (2017).

33. Liu, L. et al. One-pot cascade biotransformation for efficient synthesis of benzyl alcohol and its analogs. Chem. Asian J. 15, 1018–1021 (2020).

34. Fu, B. et al. Renewable vanillylamine synthesis from lignin-derived feedstocks. ACS Agric. Sci. Technol. 1, 566–571 (2021).

35. Yang, J. et al. Metabolic engineering of *Escherichia coli* for the biosynthesis of alpha-pinene. Biotechnol. Biofuels Bioprod. 6, 60 (2013).

36. Niu, F.X., He, X., Wu, Y.Q. & Liu, J.Z. Enhancing production of pinene in *Escherichia coli* by using a combination of tolerance, evolution, and modular co-culture engineering. Front. Microbiol. 9, 1623 (2018).

37. Ma, C., Guo, L. & Yang, H. Improved photo-hydrogen production by transposon mutant of *Rhodobacter capsulatus* with reduced pigment. Int. J. Hydrog. Energy 37, 12229–12233 (2012).

38. Ma, C., Wang, X., Guo, L., Wu, X. & Yang, H. Enhanced photo-fermentative hydrogen production by *Rhodobacter capsulatus* with pigment content manipulation. Bioresour. Technol. 118, 490–495 (2012).

39. Zheng, X., Ma, H. & Yang, H. The effect of *cycA* overexpression on hydrogen production performance of *Rhodobacter sphaeroides* HY01. Int. J. Hydrog. Energy 43, 13842–13851 (2018).

40. Ma, C., Yang, H.H., Zhang, Y. & Guo, L.J. Disruption of multidrug resistance protein gene of *Rhodobacter capsulatus* results in improved photoheterotrophic hydrogen production. Int. J. Hydrog. Energy 38, 13031–13037 (2013).

41. Zhang, Y., Yang, H.H. & Guo, L.J. Enhancing photo-fermentative hydrogen production performance of *Rhodobacter capsulatus* by disrupting methylmalonate-semialdehyde dehydrogenase gene. Int. J. Hydrog. Energy 41, 190–197 (2016).

42. Nybo, S.E., Khan, N.E., Woolston, B.M. & Curtis, W.R. Metabolic engineering in chemolithoautotrophic hosts for the production of fuels and chemicals. Metab. Eng. 30, 105–120 (2015).

43. Jodlbauer, J., Rohr, T., Spadiut, O., Mihovilovic, M.D. & Rudroff, F. Biocatalysis in green and blue: Cyanobacteria. Trends Biotechnol. 9, 875–889 (2021).

44. Sistrom, W.R. A requirement for sodium in the growth of *Rhodopseudomonas spheroides*. J. Gen. Microbiol. 22, 778–785 (1960).

45. Bose, A. & Newman, D.K. Regulation of the phototrophic iron oxidation (*pio*) genes in *Rhodopseudomonas palustris* TIE-1 is mediated by the global regulator, FixK. Mol. Microbiol. 79, 63–75 (2011).

46. Zhang, Y., Song, X., Lai, Y., Mo, Q. & Yuan, J. High-yielding terpene-based biofuel production in *Rhodobacter capsulatus*. ACS Synth. Biol. 10, 1545–1552 (2021).

47. Pal, S., Park, D.H. & Plapp, B.V. Activity of yeast alcohol dehydrogenases on benzyl alcohols and benzaldehydes: Characterization of ADH1 from *Saccharomyces carlsbergensis* and transition state analysis. Chem. Biol. Interact. 178, 16–23 (2009).

48. Yano, T., Sanders, C., Catalano, J. & Daldal, F. *sacB*-5-Fluoroorotic acid-*pyrE*based bidirectional selection for integration of unmarked alleles into the chromosome of *Rhodobacter capsulatus*. Appl. Environ. Microbiol. 71, 3014–3024 (2005).

