## Supporting information for "Harnessing photosynthetic bacterium for light-powered biocatalysis"

**Supplementary Table S1. Strains used in this study.**

| Strain | Description | Source |
|---|---|---|
| *E. coli* S17-1 | *thi pro hsdR hsdM*$^{+}$ *recA*::(RP4-2-Tc::Mu-Km::Tn7) λpir, Sm$^{r}$ | [1] |
| *R. palustris* CGA009 | Wild type | [2] |
| YC1 | CGA009 with *RPA1206* gene deletion | This study |
| YC2 | CGA009 with *RPA1206*, *RPA1687* and *RPA1725* genes deletion | This study |
| YC3 | YC2 with *ispA* promoter replaced by $P_{lac}$ | This study |
| YC4 | YC2 with both *ispA* and *crtE* promoters replaced by $P_{lac}$ | This study |
| YC5 | CGA009 with *ispA* promoter replaced by $P_{lac}$ | This study |
| YC6 | CGA009 with both *ispA* and *crtE* promoters replaced by $P_{lac}$ | This study |
| RVA1 | CGA009 harboring plasmids pBRT334O-CouBA and pGenT334O-ADH2 | This study |
| RVA2 | YC1 harboring plasmids pBRT334O-CouBA and pGenT334O-ADH2 | This study |
| RVA3/RHBA1 | YC2 harboring plasmids pBRT334O-CouA and pGenT334O-ADH2 | This study |
| RVA4/RHBA2 | YC4 harboring plasmids pBRT334O-CouA and pGenT334O-ADH2 | This study |
| PE1 | CGA009 harboring plasmids pBRT334O-ThiM-IPK-Idi and pGenT334O-GPPSps | This study |
| PE2 | YC5 harboring plasmids pBRT334O-ThiM-IPK-Idi and pGenT334O-GPPSps | This study |
| PE3 | YC6 harboring plasmids pBRT334O-ThiM-IPK-Idi and pGenT334O-GPPSps | This study |

**Supplementary Table S2. Oligonucleotides used in this study.**

| | |
|---|---|
| Ppuc-F | CG*GAATTC*TGGACGATGGTCAAATCCG |
| Ppuc-R | TT*GGTACC*TCCTCCTGAGAGACTTACG |
| Ppuf-F | CG*GAATTC*AGGGTTCTTCCGGATAGT |
| Ppuf-R | TT*GGTACC*TCACCTCCTAGTGATGG |
| PbchP-F | CG*GAATTC*TGCTCGATCGCTGGCTCG |
| PbchP-R | TT*GGTACC*CGGCTCCGTCTCCTTCCG |
| PcrtE-F | CG*GAATTC*TAACGCTTCGCTTCCGCA |
| PcrtE-R | TT*GGTACC*ACACTCCCGCGCTTCAGG |
| Pt334O-F | CG*GAATTC*TGTCTCTCTCCTGCCGTCC |
| Pt334O-R | TT*GGTACC*GGCAAAATTGTCCCTTTTCAAGTGCCTCCTTCAGATGCAA |
| Plac-F | CG*GAATTC*CGCAACGCAATTAATGTG |
| Plac-R | TT*GGTACC*GCTGTTTCCTGTGTGAAA |
| Ptac-F | CG*GAATTC*CGGTTCTGGCAAATATTC |
| Ptac-R | TT*GGTACC*TCCTGTGTGAAATTGTTA |
| Gen-F | GACAGGATGAGGATCGTTTCGCATGTTACGCAGCAGCAACG |
| Gen-R | GTCATTTCGAACCCCAGAGTCCCGCTTAGGTGGCGGTACTTGG |
| ThiM-F | TT*CGTCTC*GGATCCGAGGAGGTATATTATGCAAGTCGACCTGCTGG |
| ThiM-R | TT*CGTCTC*ACTCCTCATGCCTGCACCTCCTGC |
| IPK-F | TT*CGTCTC*AGGAGGTATATTATGGAGCTGAATATTTCCG |
| IPK-R | TT*CGTCTC*ATCCTCTACTTTGAGAATCTGATG |
| Idi-F | TT*CGTCTC*AAGGAGGTATATTATGCAAACGGAACACGTC |
| Idi-R | TT*CGTCTC*CTCGAGTTATTTAAGCTGGGTAAATGC |
| GPPSps-F1 | TT*GGTCTC*GGATCCGAGCAGAGGAGAACTAGTAT |
| GPPSps-R1 | TT*GGTCTC*ACGACACCTTGCCGTTTTCATAATA |
| GPPSps-F2 | TT*GGTCTC*TGTCGTGCGGCCATCGGATC |
| GPPSps-R2 | TT*GGTCTC*CTCGAGTCAGAGGGGGACCGACTCG |
| couA-F | TT*CGTCTC*AAGCTTAGGAGGAATAAAGTGCTCACAGGAAACGCTCA |

| | |
|---|---|
| couA-R | TT*CGTCTC*CTCGAGTCAGCTCGGCCGAACCTTGG |
| couB-F | CG*GGATCC*AGGAGGAATAAAGTGATGGACGCCATGACCGA |
| couB-R | AGAG*AAGCTT*TCACTCCAGCGCAATCACAT |
| ADH2-F | CG*GGATCC*AGGAGGAATAAAATGTCTATTCCAGAAACTCA |
| ADH2-R | AGAGA*CTCGAG*TTATTTAGAAGTGTCAACAACG |
| ispA-uF | TT*GGTACC*TCTGGAACGCTTCACCGA |
| ispA-OE-uR | ATTAATTGCGTTGCGGGAGCACTTCCGAATGGC |
| Plac-OE-iF | ATTCGGAAGTGCTCCCGCAACGCAATTAATGTG |
| Plac-OE-iR | CGGCCGAATTTTATCTGCTGTTTCCTGTGTGAAA |
| ispA-OE-dF | ACACAGGAAACAGCAGATAAAATTCGGCCGGACC |
| ispA-dR | GC*TCTAGA*CTCCTTGGCGGACAACTGCG |
| crtE-uF | GC*TCTAGA*ACCGCAATAGGTCGCATAGC |
| crtE-OE-uR | TTAATTGCGTTGCGCAGGCACAAGTGTCAGTTTA |
| Plac-OE-cF | CTGACACTTGTGCCTGCGCAACGCAATTAATGTG |
| Plac-OE-cR | CATGGCCACACTCCCGCTGTTTCCTGTGTGAAA |
| crtE-OE-dF | ACACAGGAAACAGCGGGAGTGTGGCCATGGACG |
| crtE-dR | TT*GGTACC*GGCAACCTGATAGGCTTCG |
| ALDH1-F1 | AGAG*GAGCTC*ATCTTCACCTGGCGTTCCT |
| ALDH1-OE-R1 | CTCATCACAACGATGGCCCACTCGGCTTCTTCTA |
| ALDH1-OE-F2 | AGAAGCCGAGTGGGCCATCGTTGTGATGAGGTTCA |
| ALDH1-R2 | TT*GGTACC*TGAGAATGACGACACTGCCG |
| ALDH2-F1 | GC*TCTAGA*GCCATCGTGCTGGGTGAGT |
| ALDH2-OE-R1 | TTTCATCGCATGTTCGGCGGCTGTCGAAGGTGCTG |
| ALDH2-OE-F2 | ACCTTCGACAGCCGCCGAACATGCGATGAAAGCTG |
| ALDH2-R2 | TT*GGTACC*GGGTGCCAACGAAACCAAGC |
| ALDH3-F1 | GC*TCTAGA*CTGGCGGAAGTCGGACGGGTG |
| ALDH3-OE- | ATCGTCAAGAGATCACAGGCTTTGCAGGATGGCGGT |

| | |
|---|---|
| R1 | |
| ALDH3-OE-F2 | CATCCTGCAAAGCCTGTGATCTCTTGACGATGAGCG |
| ALDH3-R2 | GG*GGTACC*TGTCGATACCGACATAGGTG |
| 16Sr-qPCR-F | GTCATCCCCACCTTCCTCGC |
| 16Sr-qPCR-R | ATGGCTGTCGTCAGCTCGTG |
| ispA-qPCR-F | GACGAAGGTGGTCTTGCCGA |
| ispA-qPCR-R | GTGAAGCGTTCCAGATTGCC |
| crtE-qPCR-F | GAAATCAATCTCGCCCATTA |
| crtE-qPCR-R | GATCTTCTCACCGACCATCC |

**Table S3. Plasmids used in this study.**

| Plasmid | Description | Source |
|---|---|---|
| pBBR1MCS-2 | Broad-host-range vector, $Km^r$ | [3] |
| pBBR-αGppsPs | pBBR1MCS-2 derivative containing GPPS-PS fusion | [4] |
| pZJD29c | Mobilizable suicide vector, $Gent^r$ | [5] |
| pBdRSf | pBBR1MCS-2 derivative removing MCS and $P_{lac}$ | [6] |
| pBRT | pBRT derivative containing the terminator rrnB T1 | This study |
| pBRPpuc | pBRT derivative carrying the promoter of *puc* operon | This study |
| pBRPpuf | pBRT derivative carrying the promoter of *puf* operon | This study |
| pBRPbchP | pBRT derivative carrying the promoter of *pgk* gene | This study |
| pBRPcrtE | pBRT derivative carrying the promoter of *eno* gene | This study |
| pBRPlac | pBRT derivative carrying the promoter of *lac* operon | This study |
| pBRPtac | pBRT derivative carrying the promoter $P_{tac}$ | This study |
| pBRPt334-6 | pBRT derivative carrying the promoter $P_{T334-6}$ of *Rhodobacter sphaeroides* | This study |
| pBRPt334O | pBRT derivative carrying the promoter of $P_{T334-6}$ and the oxygen-regulatory protein binding site of $P_{puc}$ from *R. sphaeroides* | This study |
| pGenPt334O | pBRPt334O derivative with gentamycin resistance instead of kanamycin resistance | This study |
| pBRpuc-eGFP | pBRPpuc derivative harboring the *eGFP* driven by $P_{puc}$ | This study |
| pBRpuf-eGFP | pBRPpuf derivative harboring the *eGFP* driven by $P_{puf}$ | This study |
| pBRbchP-eGFP | pBRPbchP derivative harboring the *eGFP* driven by $P_{bchP}$ | This study |
| pBRcrtE-eGFP | pBRPcrtE derivative harboring the *eGFP* driven by $P_{crtE}$ | This study |
| pBRPlac-eGFP | pBRPlac derivative harboring the *eGFP* driven by $P_{lac}$ | This study |
| pBRPtac-eGFP | pBRPtac derivative harboring the *eGFP* driven by $P_{tac}$ | This study |
| pBRPt334-6-eGFP | pBRPt334-6 derivative harboring the *eGFP* driven by $P_{T334-6}$ | This study |
| pBRPt334O-eGFP | pBRPt334O derivative harboring the *eGFP* driven by | This study |

| | | |
|---|---|---|
| | $P_{T334O}$ | |
| pBRT334O-CouBA | pBRPt334O derivative containing cascade enzymes CouB-CouA | This study |
| pGenT334O-ADH2 | pGenPt334O derivative containing enzyme ADH2 | This study |
| pBRT334O-ThiM-IPK-Idi | pBRPt334O derivative containing cascade enzymes ThiM-IPK-Idi | This study |
| pGenT334O-GPPSps | pGenPt334O derivative containing fusion enzymes GPPS/PS | This study |
| pZJ-Plac-ispA | pZJD29c derivative containing the flanking regions of *ispA* promoter and $P_{lac}$ fragment between the flanking regions | This study |
| pZJ-Plac-crtE | pZJD29c derivative containing the flanking regions of *crtE* promoter and $P_{lac}$ fragment between the flanking regions | This study |
| pZJ-ΔALDH1 | pZJD29c derivative containing the flanking regions of *RPA1206* gene | This study |
| pZJ-ΔALDH2 | pZJD29c derivative containing the flanking regions of *RPA1687* gene | This study |
| pZJ-ΔALDH3 | pZJD29c derivative containing the flanking regions of *RPA1725* gene | This study |

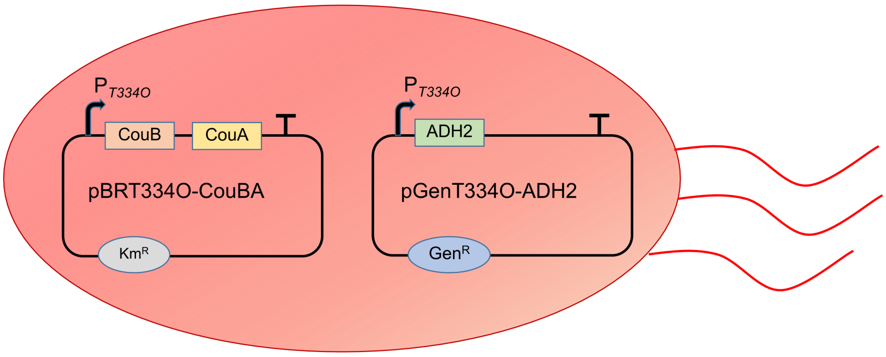


Figure S1 *R. palustris* contained two plasmids for synthesis of vanillyl alcohol (VA) or *p*-hydroxybenzyl alcohol (*p*HBA).

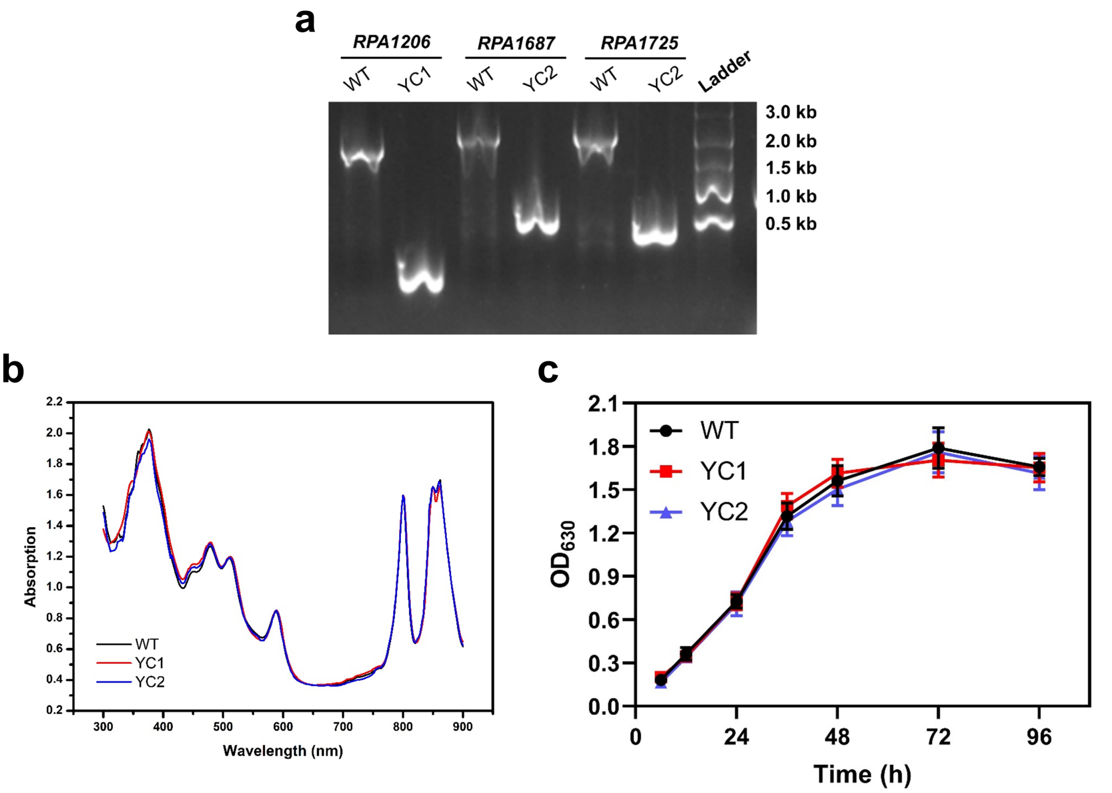


Figure S2 Effects of *aldh* deletions on *R. palustris*. a) Agarose gel image for PCR verification of genes knockout. b) The growth profiles of *R. palustris* under light-anaerobic conditions. c) The light absorption of *R. palustris* under light-anaerobic conditions. WT, YC1, and YC2 respectively represent for wild type, the mutant with *RPA1206* deletion, and the mutant with *RPA1206*, *RPA1687* and *RPA1725* deletions. Data represent the average of three replicates and error bars represent the standard deviation.

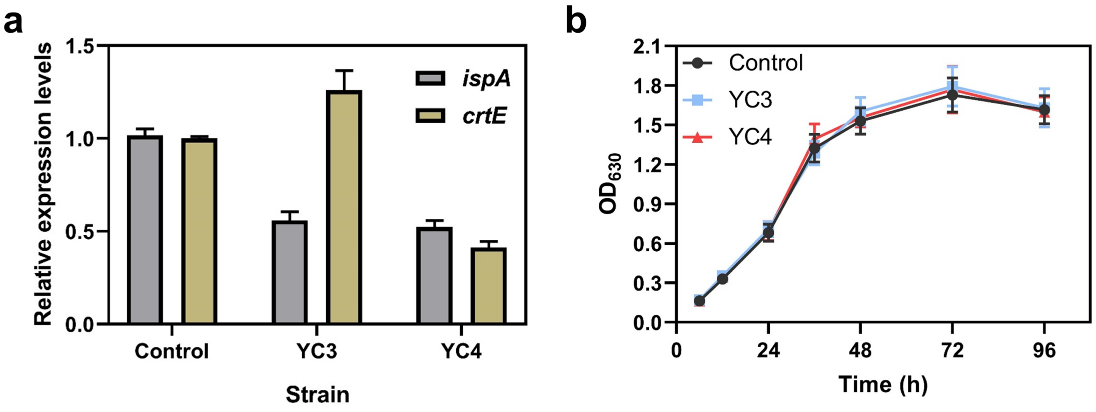


Figure S3 Effects of the decreases in *ispA* and *crtE* expressions on *R. palustris*. a) Relative expression levels of *ispA* and *crtE* genes in different strains. b) The growth profiles of *R. palustris* under light-anaerobic conditions. Control represents for strain YC2.YC3 and YC4 respectively represent for decrease in *ispA* and both *ispA* and *crtE* based on the strain YC2. Data represent the average of three replicates and error bars represent the standard deviation.

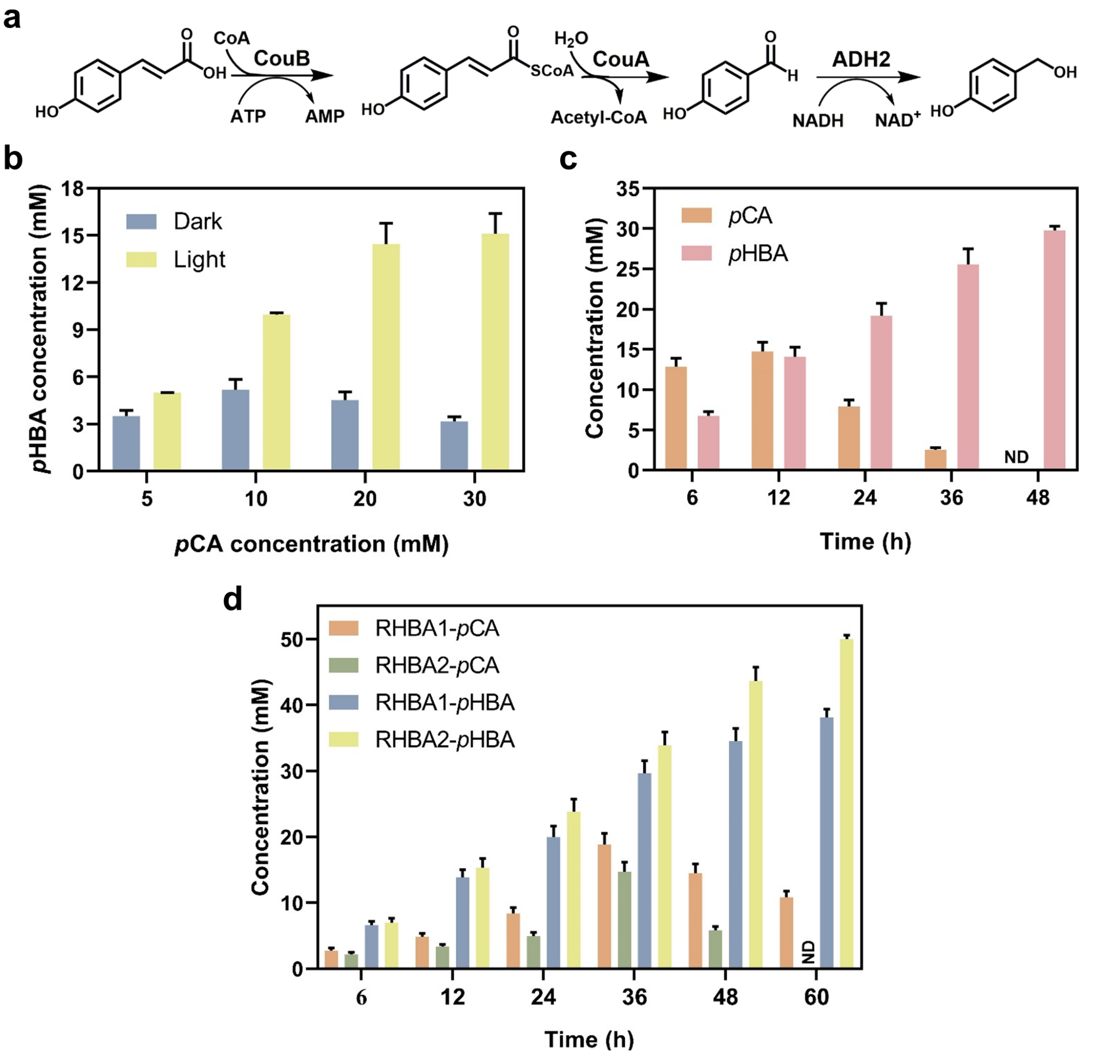


Figure S4 Synthesis of *p*HBA from *p*CA using whole-cell biocatalysis. a) Biocatalytic route for converting *p*CA into *p*HBA. b) Bioconversion of *p*CA at different concentrations into *p*HBA using the resting cells of RHBA1 under dark- and light-anaerobic conditions. c) Time course of *p*HBA production from 30 mM *p*CA by RHBA1 using the pulse-feeding approach under light-anaerobic conditions. d) *p*HBA synthesis from 50 mM *p*CA using RHBA1 and RHBA2 under the pulse-feeding mode. ND means no detection. All experiments were conducted in triplicate. Data represent the mean and standard deviations.

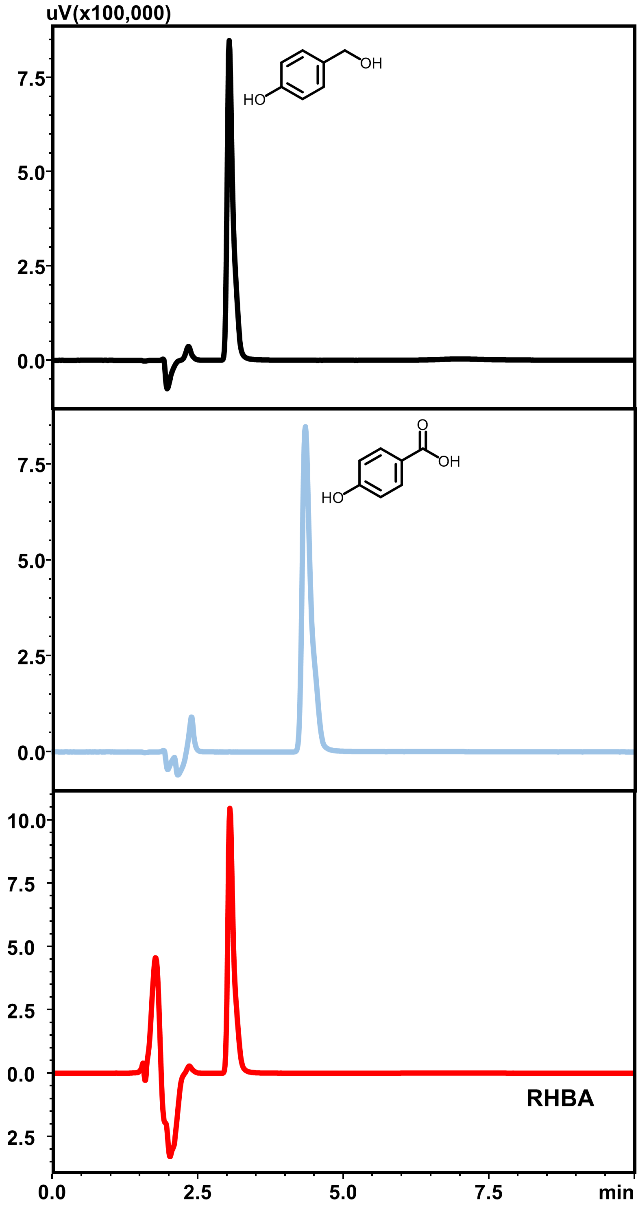


Figure S5 The HPLC result for the production of *p*HBA from *p*CA.

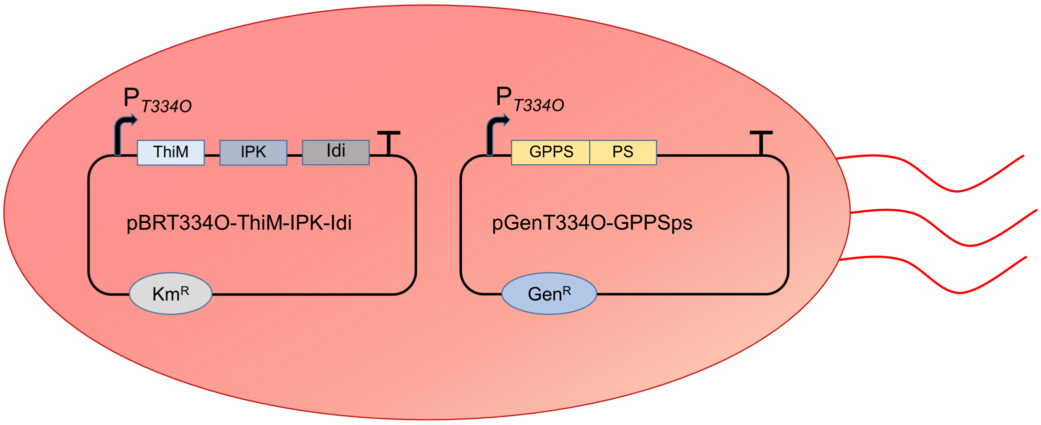


Figure S6 *R. palustris* contained two plasmids for pinene synthesis from isoprenol.

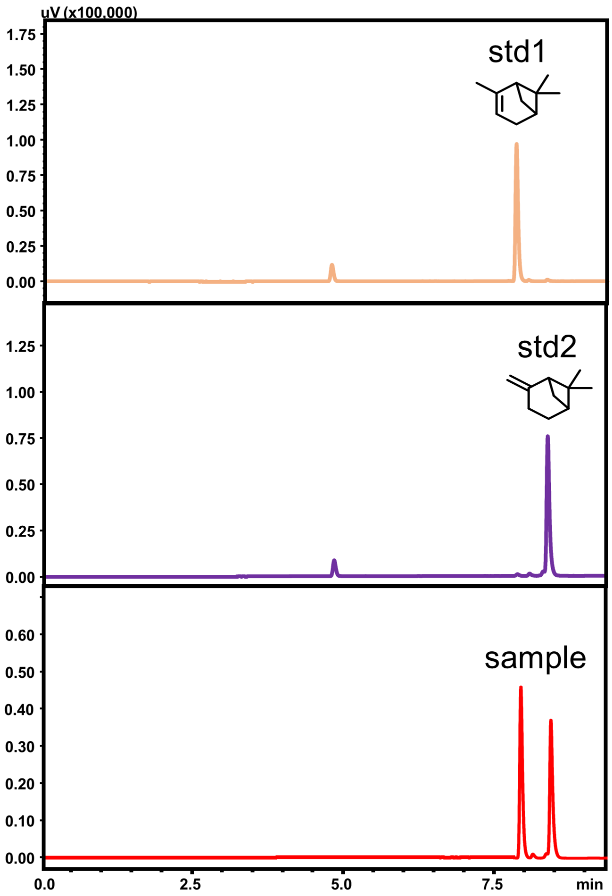


Figure S7 The GC result for the synthesis of pinene from isoprenol.

**References**:


1. Penfold, R.J. & Pemberton, J.M. An improved suicide vector for construction of chromosomal insertion mutations in bacteria. *Gene* **118**, 145-146 (1992).
2. Larimer, F.W. et al. Complete genome sequence of the metabolically versatile photosynthetic bacterium *Rhodopseudomonas palustris*. *Nat. Biotechnol.* **22**, 55-61 (2004).
3. Kovach, M.E. et al. Four new derivatives of the broad-host-range cloning vector pBBR1MCS, carrying different antibiotic-resistance cassettes. *Gene* **166**, 175-176 (1995).
4. Wu, X. et al. Biosynthesis of pinene in purple non-sulfur photosynthetic bacteria. *Microb. Cell Fact.* **20**, 101 (2021).
5. Yano, T., Sanders, C., Catalano, J. & Daldal, F. *sacB*-5-Fluoroorotic acid-*pyrE*-based bidirectional selection for integration of unmarked alleles into the chromosome of *Rhodobacter capsulatus*. *Appl. Environ. Microbiol.* **71**, 3014-3024 (2005).
6. Zhang, Y., Song, X., Lai, Y., Mo, Q. & Yuan, J. High-yielding terpene-based biofuel production in *Rhodobacter capsulatus*. *ACS Synth. Biol.* **10**, 1545-1552 (2021).
